# Transcriptome and Mutant Analysis of Neuronal Genes for Memory Formation and Retrieval in *Caenorhabditis elegans*

**DOI:** 10.1101/2023.10.31.564890

**Authors:** Yen-Ju Chen, Shyang-Jen Wu, Yu-Cheng Tang, Yu-Chun Wu, Yueh-Chen Chiang, Chun-Liang Pan

**Affiliations:** Institute of Molecular Medicine, College of Medicine, National Taiwan University, Taipei 10002, Taiwan; Center for Precision Medicine, College of Medicine, National Taiwan University, Taipei 10002, Taiwan; The Solomon H. Snyder Department of Neuroscience, Johns Hopkins University School of Medicine, Baltimore, MD 21205, USA

## Abstract

Although it is generally believed that the formation and retrieval of memory require distinct sets of genes, little is known about the identity and temporal specificity of genes that act at different memory stages. Here we characterize the transcriptomes for the formation and retrieval of stress-induced aversive memory in the nematode *Caenorhabditis elegans*. Our RNA-sequencing analysis show that neuronal genes for calcium homeostasis, membrane excitability, synaptic function and signaling are progressively activated during the formation of aversive memory. Stage-specific transcriptomes further reveal that memory formation and retrieval display distinct gene activation patterns. We carried out a candidate screen of mutants for these memory genes and verified the temporal requirement of memory function for several of them. Further in-depth characterization of *casy-1*/calsyntenin show that while the short CASY-1B isoform acts in memory retrieval, the long CASY-1A isoform functions during memory formation, and its shed extracellular N-terminus fragment is likely a critical signal for memory regulation. Our work uncovers gene expression patterns associated with distinct memory stages and provides a foundation for future mechanistic interrogation of memory functions.

## INTRODUCTION

Memory is a complex cognitive function that evolves over distinct temporal stages of formation, storage and retrieval ^1^. Anatomically, the mammalian hippocampus and insect mushroom body are critical structures for memory formation ^2, 3^, whereas distributed neural networks are believed to be sites of memory storage ^4^. While the genetic basis of memory has been intensely studied over the past three decades, few of these studies explored whether individual genes are specifically required for memory formation or retrieval. Furthermore, it remains largely unknown how the genetic structures of memory formation and retrieval differ at a global scale.

Genetically tractable model organisms, such as *C. elegans* and *Drosophila*, offer unparalleled advantages for dissecting the genetic basis of distinct memory phases ^5, 6^. In *Drosophila*, distinct genes have been partly resolved for memory formation, consolidation and retrieval ^7^. In *C. elegans*, formation and retrieval of an imprinted olfactory memory for the pathogenic bacteria *Pseudomonas aeruginosa* require distinct circuits of interneurons, but the identity of genes acting at these distinct memory stages is largely unknown ^8^. The cyclic AMP response element binding protein (CREB) is a conserved transcription factor important for memory function ^9^. But as CREB also acts in non-neural tissues to regulate a diverse range of physiological functions such as endocrine signaling and metabolism ^10^, specific CREB target genes for memory functions had been elusive until recently. Using a *C. elegans* paradigm of long-term associative memory of appetitive odors, Lakhina et al. identified memory genes regulated by CREB that showed little overlap with CREB-regulated genes for larval development and aging ^11^. Many of the mammalian and human homologs of these genes have been shown to be involved in memory functions, indicating a high degree of conservation of memory genes across species ^11^. Among these are genes encoding neurotransmitter receptors, proteins for synaptic transmission, nuclear hormone receptors and molecules for calcium signaling ^11^. A few mutants examined show defective memory retention or retrieval, whereas mutants of the *ins-22*/insulin-like peptide and *slo-1*/Big potassium (BK) channel fail to learn the association and are defective in memory formation ^11^. These observations support the use of *C. elegans* to uncover genes specifically required for memory establishment and retrieval.

Here we report neuronal gene expression patterns associated with the formation and retrieval of stress-induced aversive memory in *C. elegans*. We find that distinct genes are upregulated in either memory formation or retrieval, although a few neuronal genes are upregulated at both memory stages. We carry out a pilot candidate screen of mutant strains for these memory genes, and analyze several of them to confirm their temporal requirement for distinct memory stages. An in-depth analysis of the *casy-1*/calsyntenin mutant further reveals distinct memory functions of two CASY-1 isoforms that express in non-overlapping neurons. Our work serves as an entry point to interrogate the genetic basis of specific memory processes.

## RESULTS

### Neuronal Genes Are Progressively Upregulated during the Formation of Aversive Memory

To unravel gene expression patterns associated with the formation or retrieval of memory, we took advantage of a laboratory model of aversive associative memory triggered by mitochondrial stress ^12, 13, 14^. *C. elegans* first-stage (L1) larvae cultivated on bacteria expressing RNAi for *atp-2*/ATP synthase progressively developed avoidance of the vector bacteria over a duration of 54 h, when the worms reached young adult (Figure 1A and B). A pilot sequencing experiment was performed for bulk RNAs collected from worms subjected to *atp-2* RNAi for 24, 36 and 54 h, which corresponded to the early, middle and peak phases of bacterial avoidance (Figure 1B). Principal component analysis (PCA) showed that *atp-2* RNAi and control worms differed significantly in their gene expression patterns, with further variation contributed by the developmental stage of the animals (Figure S1). Upregulated genes at all three stages were enriched for gene ontology (GO) terms of unfolded protein responses, confirming that *atp-2* RNAi induced physiological stress (Figure 1C and 1D and Table S1). GO terms related to neuronal functions, such as chemical synapse transmission, calcium-dependent exocytosis, regulation of membrane potential and axon guidance, were enriched in the upregulated genes of *atp-2* RNAi animals at the middle and late but not early stage of stress induction (Figure 1D and Table S1), consistent with the gradual development of behavioral and memory phenotypes.

**Figure 1.**
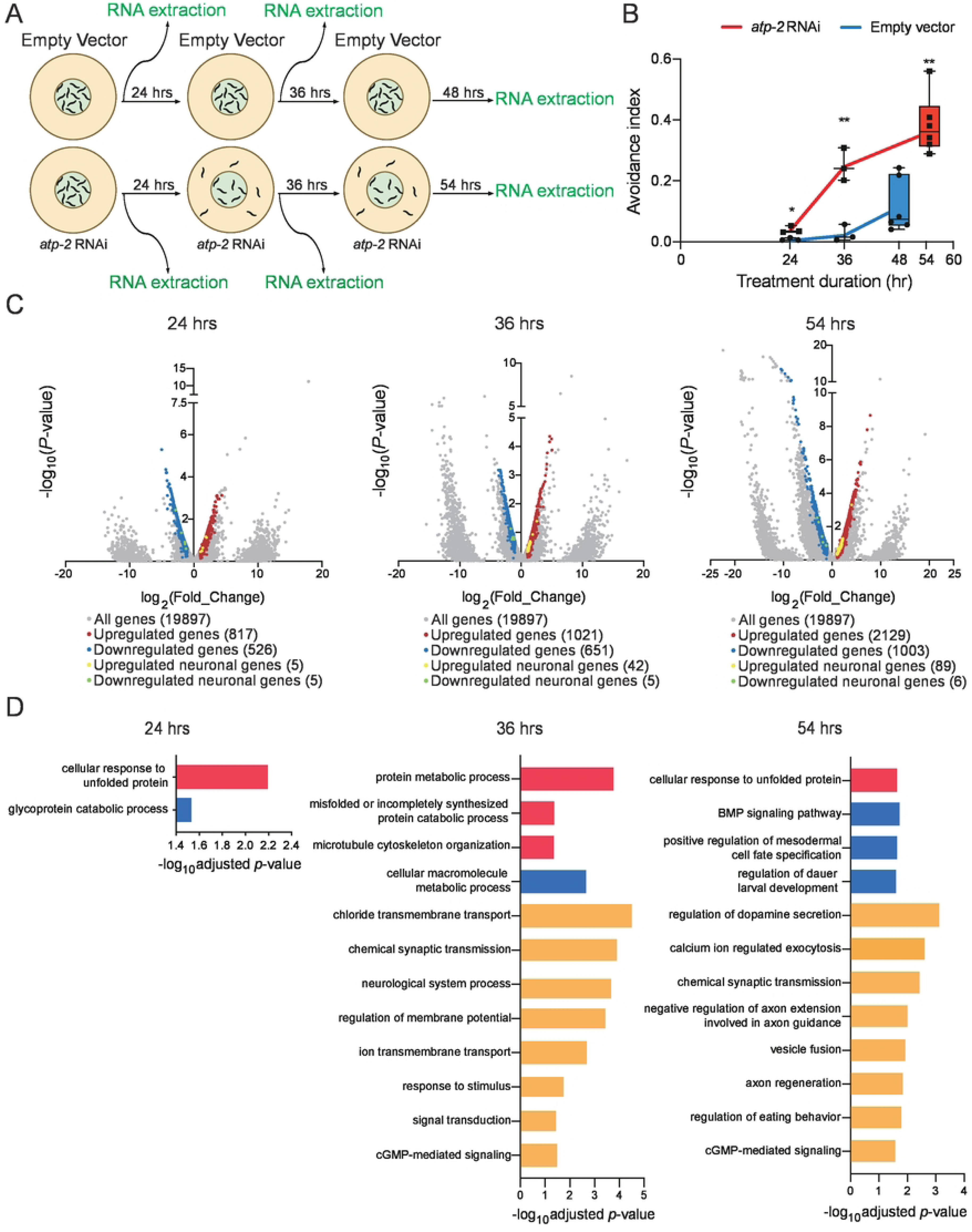
Neuronal Genes Are Progressively Activated during the Formation of Aversive Memory under Mitochondrial Stress. (A) Schematic diagram of bacterial avoidance behavior induced by *atp-2* RNAi. (B) Quantification of bacterial avoidance. Animals are treated with control and *atp-2* RNAi and their behavior is evaluated with the avoidance index before RNA is extracted at indicated time points. N = numbers of assays, with 50-200 animals per assay. Individual data points with median ± quartiles and 10 %/90 % whiskers and *p* values are indicated. Two-way ANOVA followed by Bonferroni’s correction. (C) Volcano plots of gene expression at 24, 36 and 48/54 hours after RNAi. Red, upregulated genes; blue, downregulated genes; yellow, upregulated neuronal genes; green, downregulated neuronal genes. (D) GO enrichment of upregulated genes at indicated time points of RNAi. Red and green indicate stress- and neuronal functions-related GO terms. See also Figure S1.

### Shared and Distinct Gene Expression Patterns for the Formation or Retrieval of Aversive Memory

To identify genes specifically associated with the formation or retrieval of aversive memory, we transferred *atp-2* RNAi animals to RNAi-free bacterial plates after successful induction of the avoidance behavior under stress (Figure 2A). Bacterial avoidance gradually emerged over several hours and peaked at 20 h after transfer, consistent with a prior study (Figure 2B) _12_. We designated the completion of *atp-2* RNAi (0 h) and 20 h post-transfer as the memory formation and retrieval stage, respectively. Two independent RNA samples were collected for both memory stages, followed by massive parallel sequencing. PCA showed that mitochondrial stress contributed significantly to the difference in global gene expression patterns between the *atp-2* RNAi and control animals (Figure S2). A total of 3481 and 3196 genes were upregulated during memory formation and retrieval, respectively (Figure 2C and Table S2). Of these, 2326 genes were upregulated at both stages, whereas 1155 and 870 genes were upregulated only in memory formation and retrieval, respectively (Figure 2D). GOs related to stress responses were enriched in the genes upregulated during memory formation but not retrieval, which validates our experimental scheme for memory staging (Table S3).

**Figure 2.**
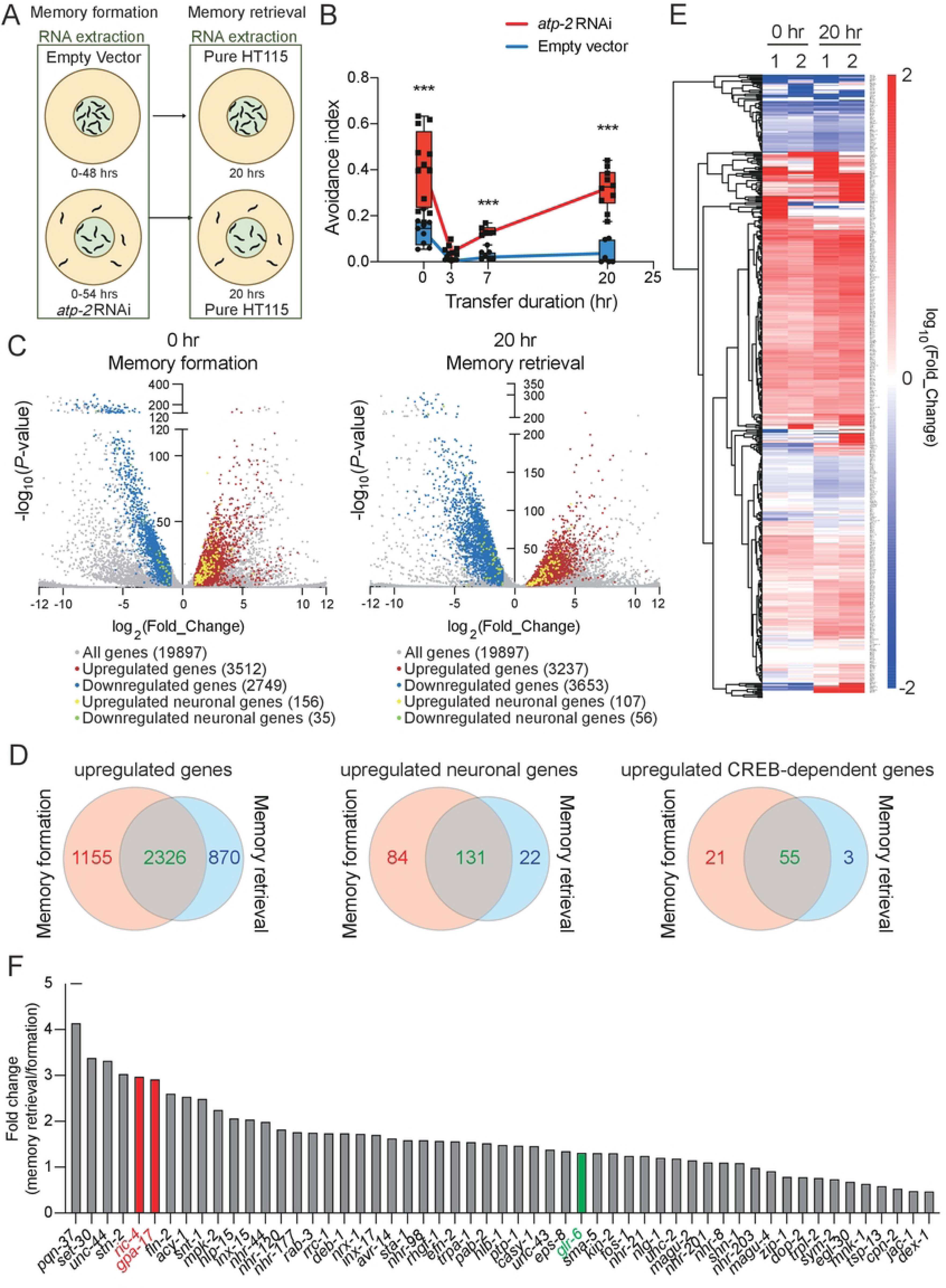
Analysis of Transcriptomes Associated with the Formation and Retrieval of Stress-Induced Aversive Memory. (A) Schematic diagram of bacterial avoidance behavior induced by *atp-2* RNAi and reactivated in the absence of RNAi. (B) Quantification of bacterial avoidance. Animals are treated with control and *atp-2* RNAi for 48-54 hours, transferred to RNAi-free plates for 20 hours, and their behavior is evaluated at indicated time points (0 h, at the end of RNAi treatment; 3 h and 20 h, 3 and 20 hours after transfer to RNAi-free plates). N = numbers of assays, with 50-200 animals per assay. Individual data points with median ± quartiles and 10 %/90 % whiskers and *p* values are indicated. Two-way ANOVA followed by Bonferroni’s correction. (C) Volcano plots of gene expression at the end of RNAi treatment and 20 hours after transfer to RNAi-free plates. Red, upregulated genes; blue, downregulated genes; yellow, upregulated neuronal genes; green, downregulated neuronal genes. (D) Cluster analysis of neuronal gene expression (n =495). (E) Venn diagrams of all upregulated genes (left), upregulated neuronal genes (middle) or CREB-dependent neuronal genes (right) between transcriptomes for memory formation and retrieval. (F) Fold change ratio between the expression levels during memory retrieval and memory formation for genes that are upregulated at both memory stages under stress compared to the control. Genes that are examined in further detail are highlighted. See also Figure S2.

A comparison of differentially expressed neuronal genes showed high consistency between independent experiments of the same memory stage (Figure 2E). Many neuronal genes displayed similar trends of differential expression between memory formation and retrieval (Figure 2E and Table S3), likely reflecting shared neural mechanisms that transduce sensory signals, integrate neural information and execute motor output to drive avoidance behaviors during both the establishment and reactivation of aversive memory. Interestingly, we found that a few neuronal genes were specifically upregulated during either memory formation or retrieval, but not both (Figure 2*D* and 2E and Table S4). A total of 131 neuronal genes were upregulated at both stages, whereas 84 and 22 neuronal genes were upregulated only during memory formation or retrieval, respectively (Figure 2*D* and 2E and Table S4). In *C. elegans*, the bZIP transcription factor CRH-1/CREB is required for aversive imprinted olfactory memory ^8^ and long-term associative memory of appetitive odors ^15^. We compared our list of differentially expressed neuronal genes to that of CREB-regulated memory genes identified in a prior *C. elegans* study, and found significant overlap between the two (Table S4) ^11^. A total of 55 CREB-regulated neuronal genes were increased at both stages, whereas 21 and 3 CREB-neuronal genes were upregulated only during memory formation or retrieval, respectively (Figure 2*D* and 2E and Table S4). We further compared the fold change of expression levels for CREB-related neuronal genes that are upregulated in both memory processes, revealing that among these genes, some are more upregulated during memory formation, and some more so during memory retrieval (Figure 2F). We conclude that CREB-regulated genes are likely conserved in multiple forms of memory with diverse temporal and contextual characteristics, and that specific CREB-neuronal genes have unique functions at distinct memory stages. We focus on these conserved CREB-regulated neuronal genes for further analysis.

### A Transcriptome-Guided Mutant Screen Validates the Importance of Memory Genes

Interestingly, the majority of the upregulated, CREB-dependent neuronal genes in our transcriptome analysis encode proteins that are implicated in the structure or function of synapses (Figure 3A). To validate the importance of the CREB-regulated neuronal genes that are increased in our memory model, we collected available single mutant strains for 45 of these genes, and subjected them to memory analysis. For these and the memory experiments in the rest of the study, instead of *atp-2* RNAi, we induced acute mitochondrial stress by using the cytochrome c reductase inhibitor antimycin A. Our recent reports showed that treatment of antimycin A for 3 to 6 h induces robust aversive memory that is indistinguishable from that induced by *atp-2* RNAi ^13, 14^.

**Figure 3.**
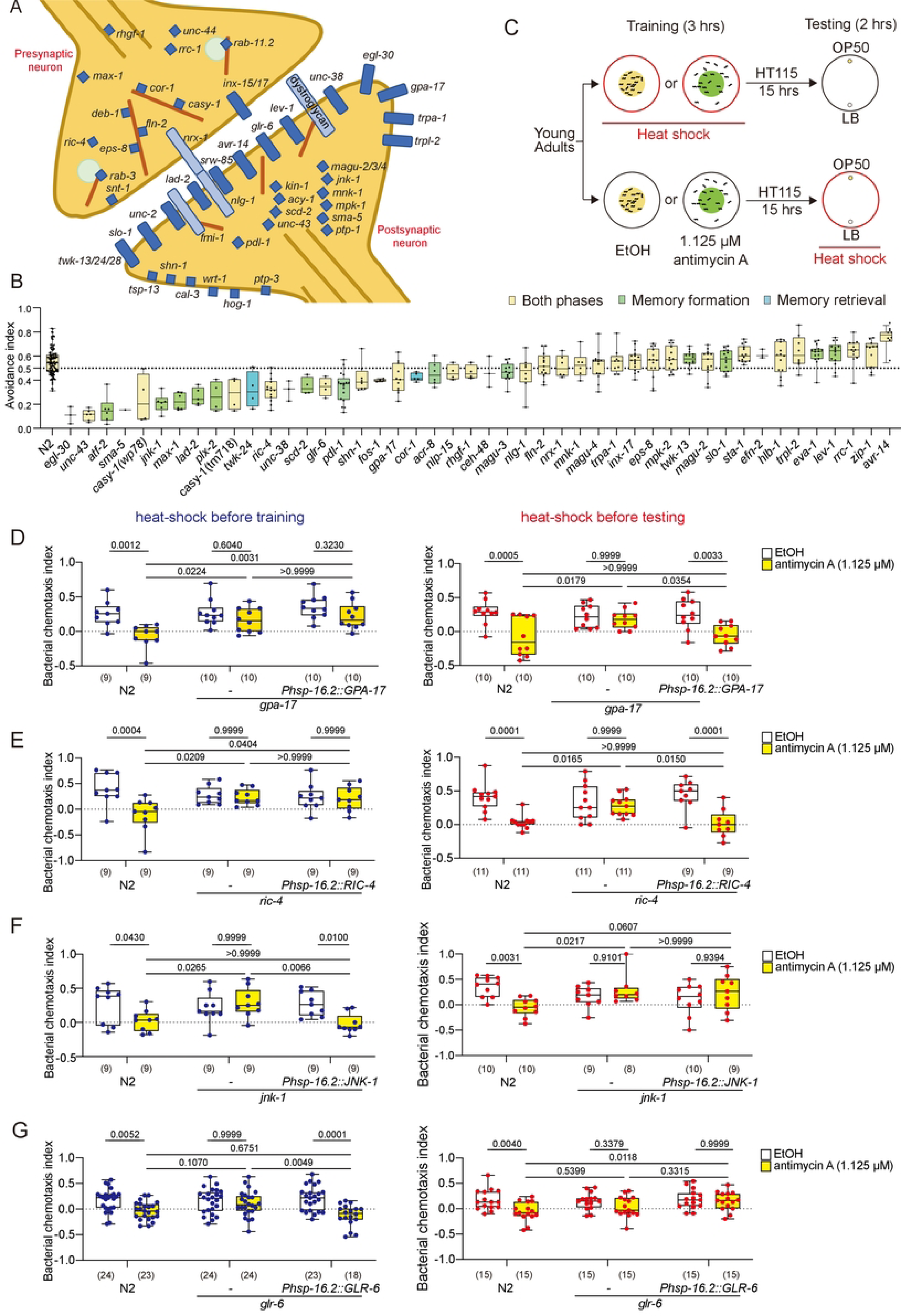
A Candidate Mutant Screen of CREB-Regulated Genes for the Formation or Retrieval of Aversive Memory. (A) Schematic diagram of upregulated CREB-dependent genes from the memory transcriptomes with respect to synaptic topography. (B) Quantification of bacterial avoidance using antimycin. N = numbers of assays, with 50-200 animals per assay. Individual data points with median ± quartiles and 10 %/90 % whiskers and *p* values are indicated. Two-way ANOVA followed by Bonferroni’s correction. (C) Schematic diagram of bacterial chemotaxis experiments coupled with heat shock-induced gene expression. (D-G) Quantification of bacterial chemotaxis index in the *gpa-17*, *ric-4*, *jnk-1*, and *glr-6* mutants with rescue by respective heat shock transgenes using the *hsp-16.2* promoter. Gene expression is induced by heat shock before training (left) or before chemotaxis (right). See Materials and Methods for details. N = numbers of assays, with 50-200 animals per assay. Individual data points with median ± quartiles and 10 %/90 % whiskers and *p* values are indicated. Two-way ANOVA followed by Bonferroni’s correction.

Compared to the wild type, about one third of these 45 mutant strains for CREB-regulated neuronal genes showed various degree of reduction in antimycin-induced bacterial avoidance (Figure 3B). These include genes for a protein tyrosine phosphatase *ptp-3*, the Gαs *gpa-17*, the L1 cell adhesion molecule (L1CAM) *lad-2*, the receptor tyrosine kinase *scd-2*, an outward rectifier potassium channel *twk-24*, the synaptic protein SHANK1 (SH3 and multiple ankyrin repeat domain 1) *shn-1* and the ionotropic glutamate receptor subunit *glr-6*. Interestingly, a mutation in the *avr-14* glutamate-gated chloride channel gene enhanced stress-induced avoidance (Figure 3B) ^16^, suggesting that some genes antagonize aversive memory. These results confirm the robustness of our stage-specific memory transcriptome analysis as an efficient screening strategy to uncover genes important for the functions of distinct memory processes.

### Examination of Genes Specifically Required for the Formation or Retrieval of Memory

We next investigate whether the temporal patterns of gene expression correlate with the requirement of gene functions for a specific memory stage. To this end, we used heat shock for stage-specific induction of gene expression (Figure 3C). We tested five genes: the Jun kinase *jnk-1,* the Gαs *gpa-17*, the SNAP25 homolog *ric-4*, the ionotropic glutamate receptor *glr-6*, and the calsyntenin *casy-1*. *jnk-1* is upregulated specifically during memory formation (Table S4). *gpa-17*, *ric-4*, *glr-6* and *casy-1* are upregulated in both memory processes. For the three genes specifically upregulated during memory retrieval, the mutants of *cor-1* and *twk-24* displayed significant developmental and locomotion defects that complicate behavioral analysis, and no mutants for *srw-85* are available. Therefore, we address the importance of retrieval-related memory genes with an alternative approach (see below). A bacterial chemotaxis assay that we recently developed was employed to test learned aversion of the food bacteria OP50 15 h after antimycin treatment ^13, 14^.

We first showed that heat-shock induction of *gpa-17* before antimycin training does not restore bacterial avoidance to the *gpa-17* mutant, while induction of *gpa-17* during the rest rescued the learned bacterial aversion of the *gpa-17* mutant (Figure 3D). These results suggest that *gpa-17* functions in memory retrieval. Similar to *gpa-17*, expressing both *ric-4a* and *ric-4b* isoforms during the rest rescued the memory deficits, while expressing them before antimycin training failed to restore the memory (Figure 3E). These results indicate that *ric-4* functions in memory retrieval, consistent with its role in mammalian memory functions ^17, 18^. With a similar approach, we showed that *glr-6* and *jnk-1* both act in memory formation instead of memory retrieval (Figure 3F and 3G). The temporal requirement of *jnk-1* matches its upregulation during memory formation (Figure 3G and Table S4). Although *gpa-17*, *ric-4* and *glr-6* are upregulated in both memory formation and retrieval, a comparison of the degree of upregulation suggests that the expression levels of *gpa-17* and *ric-4* during memory retrieval are much higher than their relative changes in memory formation (Figure 2F). By contrast, the degree of upregulation for *glr-6* was similar between memory formation and retrieval (Figure 2F). These results imply that the temporal patterns of differential neuronal gene expression could implicate gene function for specific memory processes.

### The *C. elegans* Calsyntenin CASY-1 Is Essential for Aversive Memory under Mitochondrial Stress

We next performed an in-depth analysis on *casy-1*/calsyntenin, which is upregulated during both memory formation and retrieval, and is required for stress-induced avoidance (Figure 3B and Table S4) ^19, 20, 21^. *C. elegans casy-1* regulates several forms of associative learning and memory ^22, 23^. The *casy-1* locus is predicted to encode four protein isoforms that share a transmembrane domain, an acidic region and two kinesin binding regions (KBS) (Figure 4A). The long isoform CASY-1A contains additional cadherin repeats and an LG/LNS sequence in the extracellular domain that is lacking in the short CASY-1B, CASY-1C and CASY-1D isoforms, which are almost identical. For simplicity, we will use *casy-1b*/CASY-1B to represent all three short isoforms in the rest of the study. *casy-1a* and *casy-1b* were upregulated in both memory formation and retrieval, with *casy-1a* being the predominant transcripts regardless of stress (Figure S3A and S3B). *tm718* is a 601 bp deletion of exon 4 that causes frameshift and results in reduced avoidance behavior similar to that of the null allele *wp78*, which removes the entire *casy-1* locus (Figure 4A and 4B) ^24^. The *ok739* allele is a 1304 bp deletion of intron 8 (Figure 4A), which is predicted to be a part of the promoter of the three short isoforms ^25^. All three deletion alleles of *casy-1* displayed decreased avoidance, while the phenotype of *ok739* was milder compared to that of *tm718* and *wp78*, suggesting that *ok739* is a hypomorphic allele (Figure 4B). The bacterial chemotaxis assay confirmed that *casy-1* mutants are defective in aversive associative memory (Figure 4C). Naïve *casy-1* mutants displayed intact attraction to the nutritious OP50 bacteria (Figure 4C), and their locomotion was grossly similar to that of the wild type (Figure S3C and S3D). These observations suggest that *casy-1* plays a specific role in associative memory.

**Figure 4.**
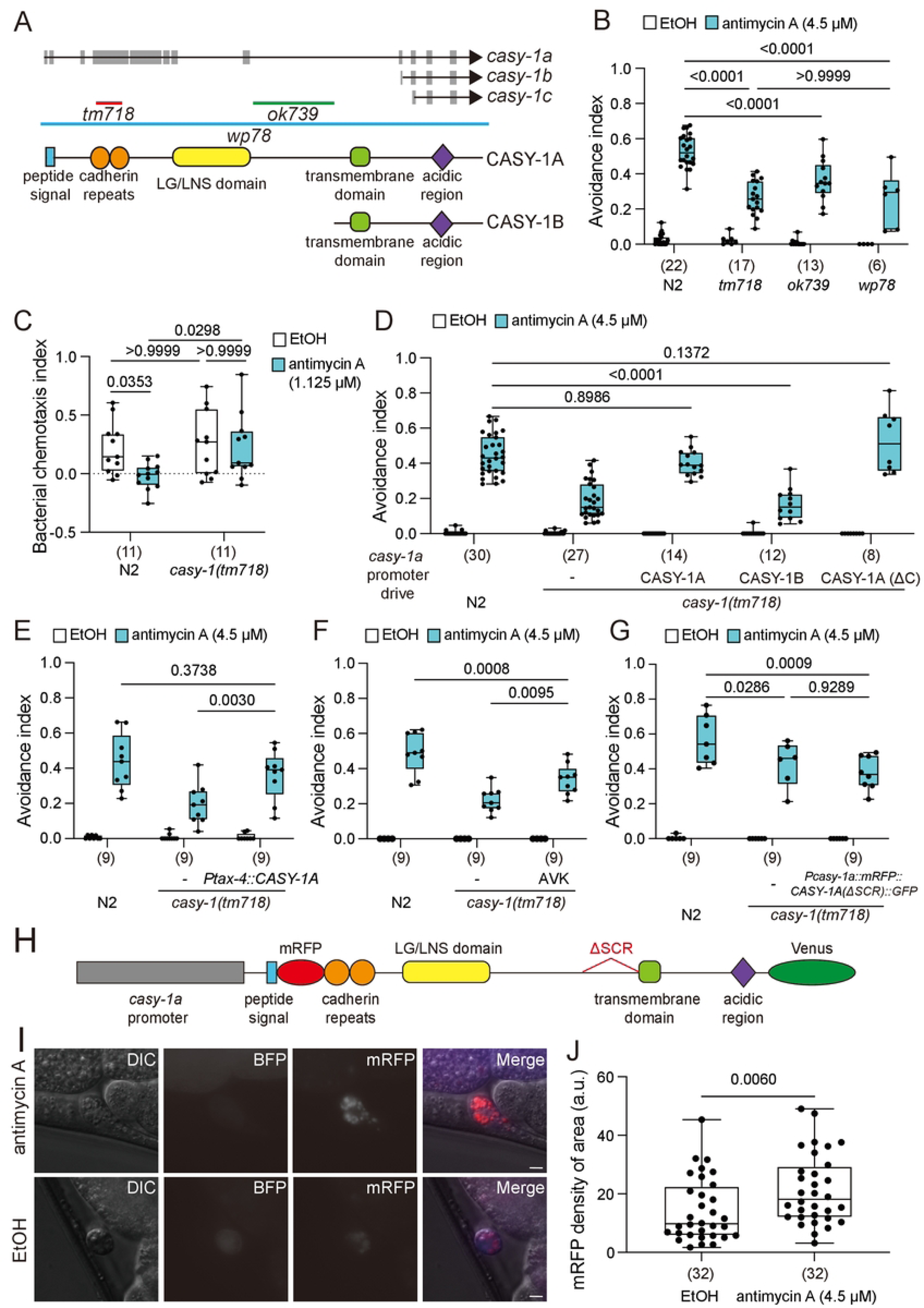
The *C. elegans* Calsyntenin CASY-1 Is Essential for Aversive Memory under Mitochondrial Stress. (A) Schematic diagrams of the *casy-1* gene and two protein isoforms. Grey boxes and black lines are exons and introns, respectively. (B, C) Quantification of bacterial avoidance in different *casy-1* alleles using the standard assay (B) or the transfer assay (C). (D-G) Quantification of bacterial avoidance in the *casy-1(tm718)* mutant expressing different *casy-1* cDNA variants from the *casy-1a* promoter (D, G) or wild-type *casy-1a* from specific neuronal promoters (E, F). N = numbers of assays, with 50-200 animals per assay. Individual data points with median ± quartiles and 10 %/90 % whiskers and *p* values are indicated. Two-way ANOVA followed by Bonferroni’s correction. (H) Schematic diagram of the CASY-1A protein tagged with mRFP and Venus at the N- and C-terminus, respectively. ΔSCR, deletion of the cleavage sequence by α- or γ-secretases (70 amino acids). (I) Epifluorescent images of the coelomocytes, which are labeled by *Punc-122::BFP*, in animals with *Pcasy-1a::mFRP::CASY-1A::Venus.* Scale bar = 5 μm. (J) Quantification of mRFP signal intensity in the coelomocytes. n = numbers of coelomocytes scored. Individual data points with median ± quartiles and 10 %/90 % whiskers are indicated. Mann-Whitney U test. See also Figures S3 and S4.

### CASY-1A Acts in Sensory Neurons and Is Shed During Memory Formation

A previous study showed that the long CASY-1A and short CASY-1B isoforms display distinct, largely non-overlapping expression patterns ^25^. We confirm this finding, showing that the *casy-1a* promoter was strongly expressed in many head and tail neurons, while the *casy-1b* promoter was active mainly in the motor neurons of the ventral nerve cord and a few head neurons (Figure S3E). Expression of CASY-1A, but not CASY-1B, from the *casy-1a* promoter completely rescued the memory defects (Figure 4D), suggesting that in *casy-1a*-expressing cells, the extracellular domains of CASY-1 are essential for its memory function. Furthermore, a truncated CASY-1A that lacks the intracellular domains also rescued the memory deficits of the *casy-1(tm718)* mutant (Figure 4D), suggesting that in CASY-1A(+) neurons, membrane-tethered N-terminal ectodomains of CASY-1A are sufficient for memory regulation. Expression of CASY-1A in 10 pairs of head sensory neurons from the promoter of *tax-4*/cyclic nucleotide-gated calcium channel restored bacterial avoidance to the *casy-1* mutant (Figure 4E). Further cell-specific rescue experiments showed that *casy-1a* has some functions in AVK but does not act in ASEL, ASI, NSM, RIA, RIC, or *glr-1*/AMPA receptor-(+) interneurons (Figures 4F and S4A). These results suggest that CASY-1A act in head sensory neurons but do not rule out its roles in other untested interneurons.

One of the established functions of CASY-1A in *C. elegans* is inter-cellular adhesion ^26^. Our structure-function analysis of CASY-1A also suggests that the ectodomains of CASY-1A are important for associative memory under stress. A previous study showed that the N-terminus extracellular fragment of CASY-1A was released during aversive salt memory ^22^. Supporting this observation, we found that CASY-1A that lacks the potential cleavage sequences for α- or γ-secretase (CASY-1A(ΔSCR)) failed to rescue the behavioral phenotype of the *casy-1* mutant (Fig. 4*G* and *H*). To test whether the CASY-1A N-terminus fragment is shed during the establishment of stress-induced aversive memory, we expressed a functional CASY-1A dually tagged with mRFP and Venus at the N- and C-terminus, respectively (Figure 4H) ^22^. The cleaved N-terminal fragment of CASY-1A is released into the pseudocoelom and endocytosed by the coelomocytes, which are constitutive phagocytic cells whose endocytosed contents serve as a proxy for extracellular secretion from other cells (Figure 4I and 4J). Deletion of the proteolytic cleavage sequence prevented CASY-1A shedding, evidenced by the absence of coelomocyte mRFP signals with reciprocal increase of mRFP signals in the source neurons (Figure S4B-E). Antimycin treatment increased mRFP signals in the coelomocytes of animals cultivated on bacterial food, suggesting increased shedding of the CASY-1A N-terminus fragment (Figure 4I and 4J). Food or antimycin alone did not increase CASY-1A shedding (Figure S4F and S4G), implying that shed CASY-1A N-terminus fragment does not simply signal food sensation or stress. Bacterial odors alone increased CASY-1A shedding in the presence of mitochondrial stress (Figure S4H), which is consistent with CASY-1A acting in sensory neurons (Figure 4E), and suggests that CASY-1A is likely a signal that associates perceived sensory cues with systemic stress.

### CASY-1B Acts in GABAergic Neurons

As CASY-1A and CASY-1B show almost mutually exclusive expression patterns (Figure S3E), we wonder whether CASY-1B also plays a role in stress-induced aversive memory. Interestingly, both CASY-1B and CASY-1A fully rescued the memory deficits of the *casy-1(tm718)* null mutant when expressed from the *casy-1b* promotor (Figure 5A). CASY-1B is expressed in the ventral cord motor neurons that are either cholinergic or GABAergic ^25^. CASY-1B expressed in the GABAergic motor neurons, using the *unc-25* promoter, was sufficient to rescue the memory phenotypes (Figure 5B). Expression of CASY-1B in the RIS GABAergic interneuron from the *flp-11* promoter partially restored bacterial avoidance, suggesting that CASY-1B functions in RIS and other GABAergic neurons (Figure 5C). By contrast, CASY-1B expression in the cholinergic motor neurons, using the *unc-17* promoter, failed to rescue (Figure 5B). These results suggest that CASY-1B functions in specific GABAergic neurons, and that when overexpressed, CASY-1A can substitute for CASY-1B in memory function.

**Figure 5.**
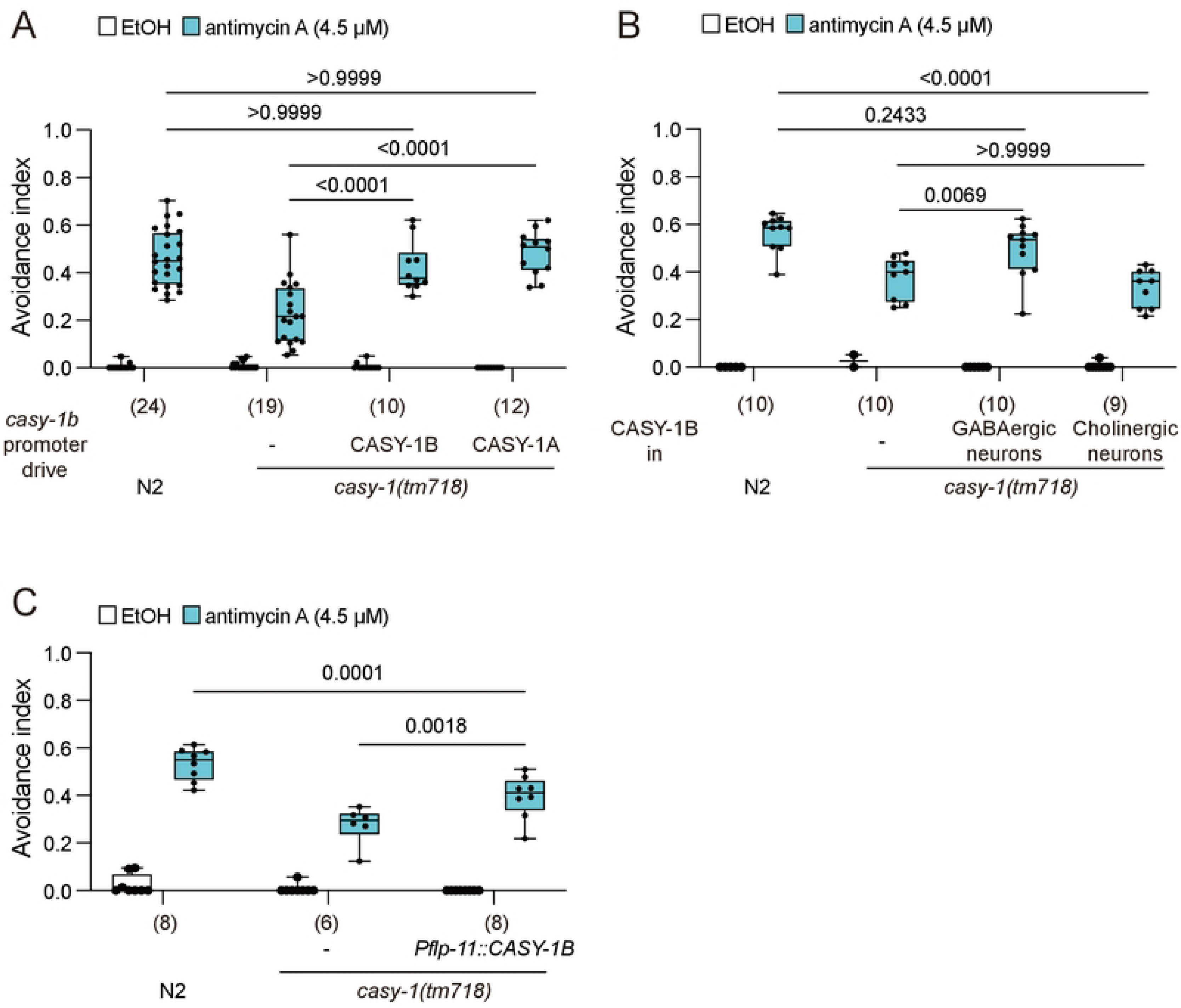
CASY-1B Functions in GABAergic Neurons, including RIS. (A) Quantification of bacterial avoidance in the *casy-1* mutant with either CASY-1 isoform from the *casy-1b* promoter (A), with CASY-1B in GABAergic (the *unc-25* promoter) or cholinergic neurons (the *unc-17* promoter) (B), or with CASY-1B in the GABAergic RIS interneurons using the *flp-11* promoter (C). N = numbers of assays, with 50-200 animals per assay. Individual data points with median ± quartiles and 10 %/90 % whiskers and *p* values are indicated. Two-way ANOVA followed by Bonferroni’s correction.

### CASY-1A and CASY-1B Control Memory Formation and Retrieval, Respectively

*casy-1* is upregulated during both memory formation and retrieval (Figure S3A and Table S4). To explore the specific memory process that *casy-1* acts in, we performed heat shock rescue experiments mentioned earlier in this study (Figure 3C). Heat-shock induction of CASY-1A before training, but not in the testing phase, significantly increased bacterial avoidance of the *casy-1* mutant (Figure 6A and 6B). This implies that CASY-1A is required for memory formation. In contrast, CASY-1B expression specifically during rest before bacterial chemotaxis, but not before training, partially rescued bacterial avoidance, suggesting that CASY-1B functions in memory retrieval (Figure 6C and 6D). These results reveal distinct functions of CASY-1 isoforms for specific memory processes.

**Figure 6.**
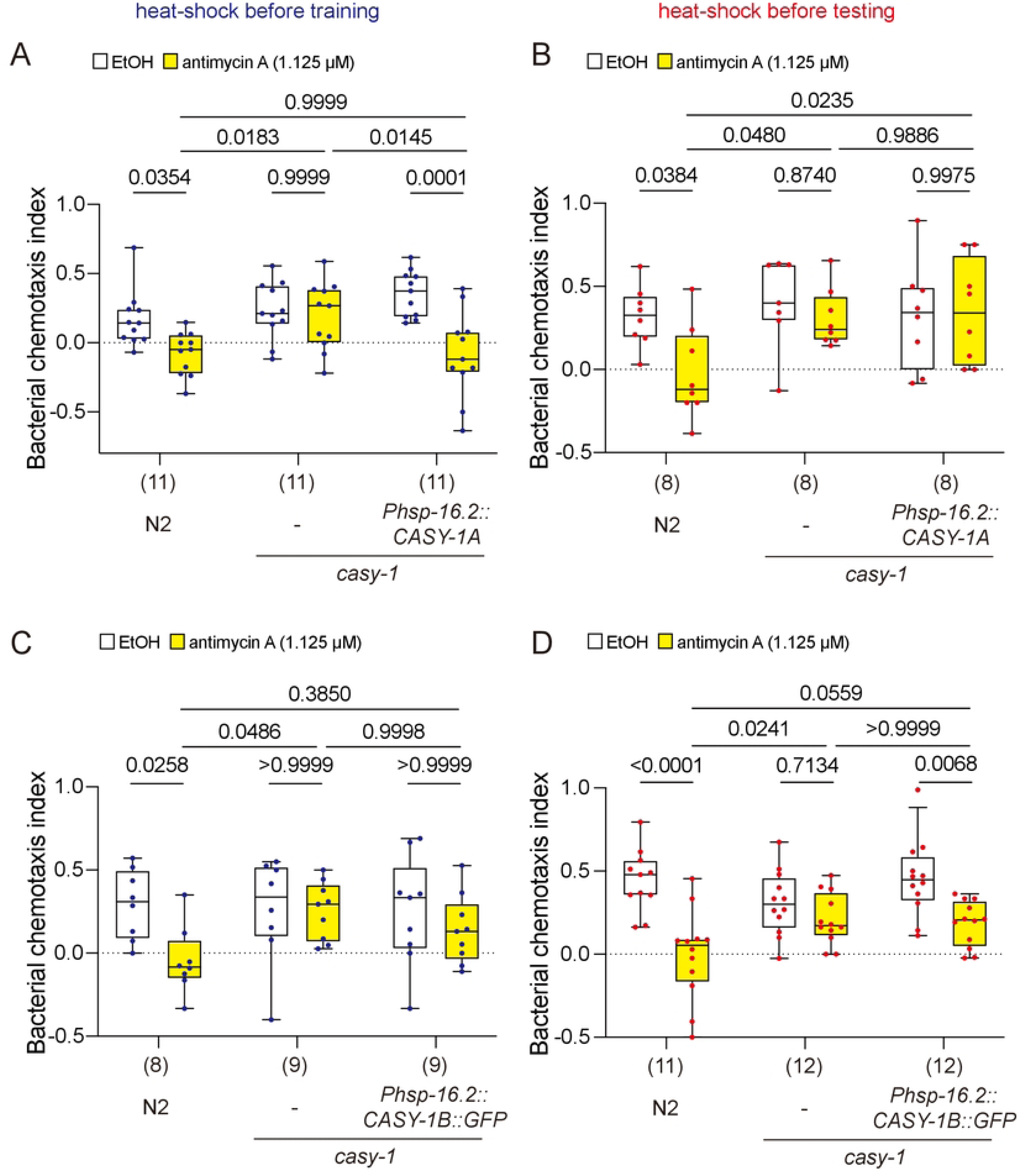
CASY-1A and CASY-1B Regulate Memory Formation and Retrieval, Respectively. (A-D) Bacterial chemotaxis index of the *casy-1* mutant expressing *Phsp-16.2::CASY-1A* (A, B) or *Phsp-16.2::CASY-1B::GFP* (C, D). CASY-1A or CASY-1B proteins are induced by heat shock at 34οC before training (A, C) or before chemotaxis (B, D). See Materials and Methods for details. N = numbers of assays, with 50-200 animals per assay. Individual data points with median ± quartiles and 10 %/90 % whiskers and *p* values are indicated. Two-way ANOVA followed by Bonferroni’s correction.

### CASY-1 Acts with the Neurexin-Like BAM-2 to Regulate Aversive Memory under Stress

Finally, we explored genes that interact with *casy-1* to promote stress-induced aversive memory. CASY-1 transports the insulin-like neuropeptide receptor DAF-2c to the presynaptic region of the ASER gustatory neuron to drive aversive salt memory under starvation ^27^. Bacterial avoidance under mitochondrial stress was decreased in both the *casy-1* and the *daf-2* mutants, and the *casy-1; daf-2* double mutant showed a significantly lower avoidance than either single mutant, suggesting that *casy-1* and *daf-2* function in parallel (Figure 7A). RIG-6 is the sole Contactin adhesion molecule in *C. elegans* and has been shown to be important for axon guidance ^28^. The modest reduction in bacterial avoidance of the *rig-6* mutant was further enhanced by the *casy-1* mutation (Figure 7B), indicating that *casy-1* and *rig-6* act independently. Vertebrate calsyntenin-3 is known to physically interact with neurexin, a conserved adhesion molecule important for synaptic functions ^29, 30^. Mutations of *nrx-1*, the *C. elegans* homolog of neurexin, neither affected bacterial avoidance on its own, nor did it enhance the avoidance defects of the *casy-1* mutant, implying that *nrx-1* is dispensable (Figure 7C). This is consistent with a lack of physical interaction between CASY-1 and NRX-1 in *C. elegans* ^26^. CASY-1 physically interacts with BAM-2, a neurexin-related transmembrane protein ^31^, and they promote axon bundling in *C. elegans* nervous system development ^26^. A mutation of *bam-2* reduced bacterial avoidance under mitochondrial stress, and it did not further decrease the low bacterial avoidance of the *casy-1* mutant (Figure 7D). This result suggests that *bam-2* and *casy-1* act in a common genetic pathway for stress-induced aversive memory. Expression of a *bam-2* cDNA in cholinergic neurons restored avoidance to the *bam-2* mutant (Figure 7E), whereas expression in the dopaminergic neurons, using the *dat-1* promoter, failed to rescue (Figure 7F). We speculate that BAM-2 functions as a receptor for the N-terminus fragment of CASY-1A to transduce signals required to form aversive memory under mitochondrial stress.

**Figure 7.**
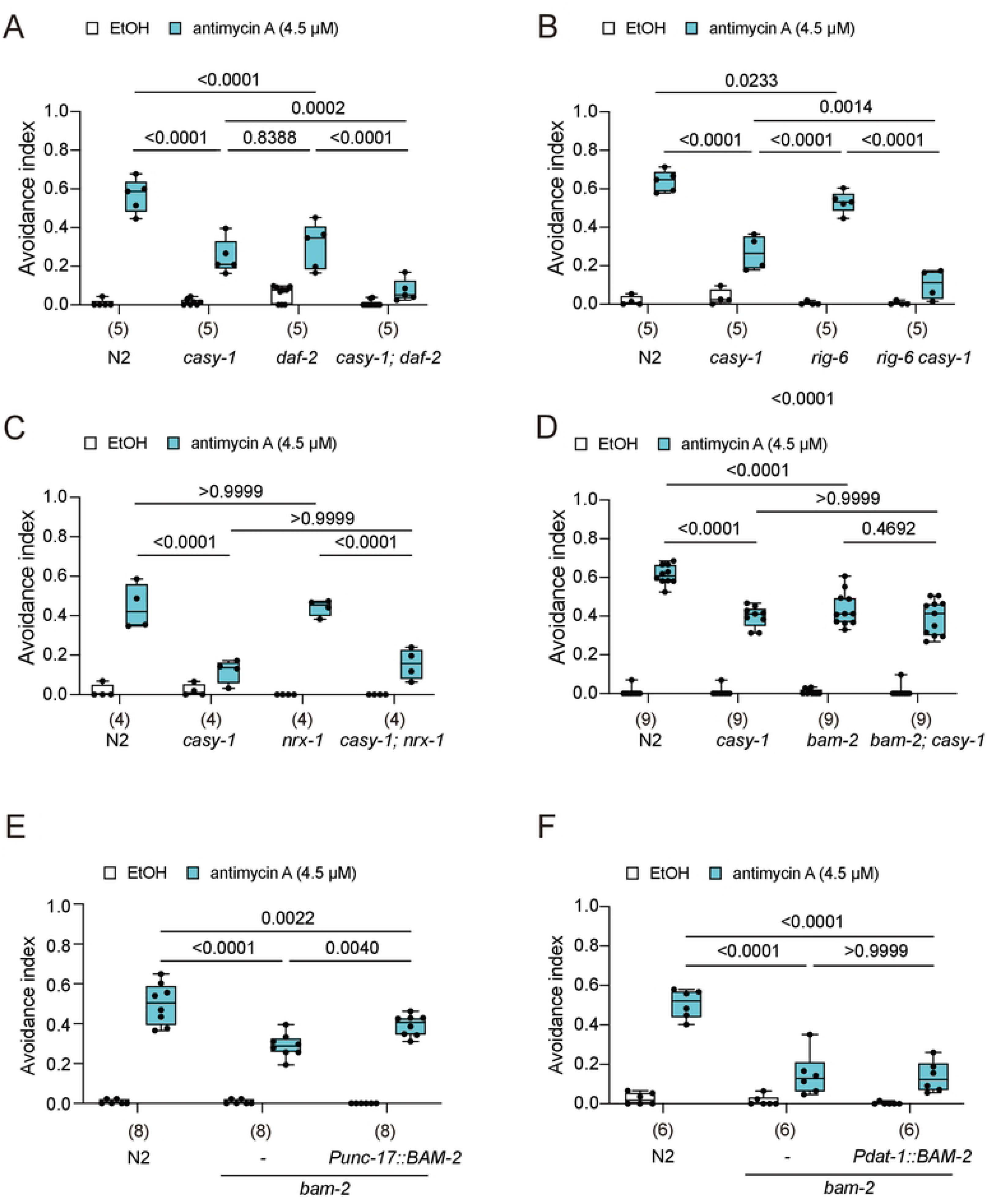
CASY-1 Acts with the Neurexin-Like Molecule BAM-2 to Regulate Aversive Memory under Stress. (A-F) Quantification of bacterial avoidance in the wild type, *casy-1(tm718)* and *daf-2(e1370)* (A), *rig-6(ok1589)* (B), *nrx-1(ok1649)* (C), *bam-2(cy6)* (D), and *bam-2* rescue in the cholinergic neurons (E, from the *unc-17* promoter) or in the dopaminergic neurons (F, from the *dat-1* promoter). N = numbers of assays, with 50-200 animals per assay. Individual data points with median ± quartiles and 10 %/90 % whiskers and *p* values are indicated. Two-way ANOVA followed by Bonferroni’s correction.

## DISCUSSION

Unsolved mysteries of memory include the anatomical substrates and genetic basis of distinct memory processes, such as memory formation and retrieval. By combining stage-specific RNA-sequencing and mutant analysis, we uncovered neuronal genes that act specifically in the establishment or recall of a stress-induced aversive memory in *C. elegans*. Two prior *C. elegans* studies profiled gene expression patterns for long-term associative memory (LTAM) by microarray analysis ^11, 32^. Using next generation RNA sequencing, our study confirms and extends these earlier findings. Even with major differences in the methodology of gene expression profiling, our catalogue of differentially expressed genes overlaps significantly with those identified in Lakhina et al. ^11^. With the advancement of gene sequencing technology, it should be possible to resolve memory gene expression at single-neuron resolution and finer temporal scales in near future.

### Shared and Unique Genes for the Formation and Retrieval of Aversive Associative Memory

Although it is widely accepted that the formation, storage and retrieval of memory occur at distinct anatomical sites in the brain, it remains largely obscure whether these memory processes also require unique sets of genes. By using a behavioral paradigm that separates distinct memory processes, we uncovered differentially expressed genes for the formation and retrieval of memory, respectively. A collection of neuronal genes is shared by both memory processes, many previously shown to be regulated by CREB ^11^, suggesting these genes are central to all memory stages. In addition, a handful of neuronal genes are specifically upregulated in only memory formation or retrieval, but not both. Past memory studies mostly focus on single genes, while large-scale transcriptomics or proteomics studies seldom provide in-depth investigation of individual neuronal genes. To circumvent these problems, we use our transcriptome data to guide a mutant screen for many candidates in the list of upregulated, CREB-related neuronal genes. For the 55 CREB-regulated neuronal genes whose expression is increased in both memory formation and retrieval, we examined 28 and found 6 were symptomatic when mutated (*casy-1*/calsyntenin, *ptp-3*/protein tyrosine phosphatase, *ric-4*/SNAP-25, *shn-1*/SHANK1, *gpa-17*/Gα, *glr-6*/ ionotropic glutamate receptor). Of the 21 CREB-related neuronal genes upregulated during memory formation, we tested 15 and found 7 to be important (*atf-2*/NFIL3 transcription factor, *jnk-1*/Jun kinase, *lad-2*/L1CAM, *max-1*/PH/MyTH4/FERM domain protein, *pdl-1*/phosphodiesterase 6D, *plx-2*/plexin and *scd-2*/ALK). Finally, of the three CREB-related neuronal genes specifically upregulated during memory retrieval, two seemed to be important (*cor-1*/Coronin, *twk-24*/potassium channel). A role in memory function has not been shown for many of these genes in the past. Although in-depth analysis of memory phenotypes is necessary for most of these genes, our combined approach of transcriptomic analysis and candidate screen proves to be promising, and results of the current study are a valuable resource for future memory research.

To corroborate the temporal patterns of gene upregulation with functional implications, we further tested *jnk-1, gpa-17*, *ric-4*, *glr-6* and *casy-1* for their requirement in specific memory processes by heat shock induction. *jnk-1* is upregulated and acts in memory formation. *glr-6*, *gpa-17* and *ric-4* are upregulated in both memory formation and retrieval. Upregulation of *gpa-17* and *ric-4* is significantly more pronounced in memory retrieval, and they also function to facilitate memory recall. By contrast, upregulation of *glr-6* is not obviously biased towards either memory stage, and rescue experiments suggest that it acts in memory formation. These observations suggest that temporal patterns of differential gene expression and gene functions are correlated to a certain degree. Further transcriptome studies are necessary to confirm and extend our findings.

### Dual Functions of CASY-1/Calsyntenin in Aversive Associative Memory

Calsyntenins are synaptic calcium-sensitive proteins that undergo proteolytic processing ^21^. They are shown to regulate cognitive functions, and in *C. elegans*, CASY-1 is important for associative learning and memory ^22, 27^. In aversive salt learning under starvation, *casy-1* regulates axonal expression of the DAF-2c isoform of the insulin-like growth factor receptor in a salt-sensing neuron ^27^. It had been shown that the intracellular domains of CASY-1 are dispensable for aversive salt learning ^22^, and that the CASY-1 ectodomains are shed likely through proteolytic cleavage ^22^. In the current study, we extend these observations and show that *casy-1* regulates aversive memory under systemic mitochondrial stress. We further provide evidence that proteolytic cleavage of CASY-1A is critical for its memory functions, as removal of the putative α- or γ-secretase sequence abolishes shedding and CASY-1A memory function. Calsyntenins, the mammalian homologs of CASY-1, undergo proteolytic cleavage by α- or γ-secretases ^33, 34^. Interestingly, the long and short isoforms of *casy-1* act in non-overlapping neurons and have distinct roles in memory processes. The long isoform CASY-1A acts in the sensory neurons, shed the ectodomains and genetically interact with the neurexin-like adhesion molecule BAM-2. In agreement with prior studies ^22^, the intracellular domains of CASY-1A are dispensable. Unexpectedly, the short isoform CASY-1B, which lacks most of the ectodomains, can also regulate memory from the GABAergic neurons when overexpressed, and it functions during memory retrieval but not memory formation. As the expression levels of *casy-1b* and *casy-1c* are very low at baseline or under stress (*SI Appendix* Fig. S3*A*), the significance of CASY-1B rescue should be cautioned, as it is likely due to overexpression of the multi-copy gene array. Interestingly, increase in CASY-1A ectodomain shedding requires both sensory cues and systemic mitochondrial stress, raising an intriguing possibility that shed CASY-1A ectodomains serve as signals for associating extrinsic stimuli with the internal physiological state. As BAM-2 genetically interacts with CASY-1, we speculate that CASY-1A relays memory information from the sensory neurons to BAM-2-expressing neurons. It remains to be determined in which neurons BAM-2 acts, as well as the signal transduction pathways downstream of BAM-2. With temporally resolved gene expression profiles coupled with a pilot mutant screen, our work serves as an entry point for future interrogation of neuronal gene functions in distinct memory processes.

## ACKNOWLEDGEMENTS

We thank Scott Emmons, Yuichi Iino and Peri Kurshan for *C. elegans* strains, and Yi-Ting Tsai, Hsin-Yue Tsai and Yen-Ping Hsueh for advice on transcriptome analysis. Some of the strains used in this study are provided by the *Caenorhabditis* Genetics Center, which is supported by NIH Office of Research Infrastructure Programs (P40OD010440), and by the National BioResources Project (NBRP), which is supported by the Japanese government. Gene database and bioinformatics analysis are provided by WormBase, which is supported by grant #U24 HG002223 from the National Human Genome Research Institute at the US National Institutes of Health, the UK Medical Research Council and the UK Biotechnology and Biological Sciences Research Council. This study was funded by the National Science and Technology Council, and the Featured Areas Research Center Program within the framework of the Higher Education Sprout Project by the Ministry of Education (MOE), Taiwan, to C.-L.P. (MOE 111L901402A, MOE 112L901402A, NSTC 110-2634-F-002-017, NSTC 110-2320-B-002-055-MY3, NSTC 112-2320-B-002-018-MY3).

## AUTHOR CONTRIBUTIONS

Y.-J.C, S.-J.W, and C.-L.P designed research; Y.-J.C, S.-J.W, Y.-C.T, Y.-C.W and Y.-C.C performed research; Y.-J.C, S.-J.W, Y.-C.T, Y.-C.W, Y.-C.C and C.-L.P analyzed data; Y.-J.C, S.-J.W and C.-L.P wrote the paper with contributions from Y.- C.T, Y.-C.W and Y.-C.C.

## INCLUSION AND DIVERSITY STATEMENT

The author list of this paper includes contributors from the location where the research was conducted who participated in the data collection, design, analysis, and /or interpretation of the work.

## STAR METHODS

## RESOURCE AVAILABILITY

### Lead Contact

Further information and requests for strains, reagents and protocols should be directed to and will be fulfilled by the lead contact, Chun-Liang Pan

### Materials Availability

*C. elegans* strains and DNA constructs generated in this study are available from the Lead Contact upon request.

### Data and Code Availability

- All the primary data used in this study have been deposited at Figshare and are publicly available as of the date of publication. The DOI is listed in the key resources table. Microscopy data reported in this study will be shared by the lead contact upon request.
- This study does not use original code for data analysis.
- Any additional information required to reanalyze the data reported in this paper is available from the lead contact upon request.

## EXPERIMENTAL MODEL AND SUBJECT DETAILS

### Animals

### *C. elegans* and *E. coli* Bacterial Strains

*C. elegans* strains are cultivated and maintained on solid nematode growth medium (NGM) seeded with OP50 or HT115 *E. coli* strains at 20°C as previously described ^35^. A list of mutant alleles and transgenic strains used in this study are listed in KEY RESOURCES TABLE.

### Microbe Strains

The OP50 and HT115 *E. coli* strains are used.

## METHODS DETAILS

### Molecular Biology and Germline Transformation

Standard molecular biology techniques were used in this study. Promoter sequences used included *hsp-16.2* (263 bp), *ric-4* (2.6 kb), *casy-1a* (2.0 kb), *casy-1b* (3.1kb), *tax-4* (4.5 kb*), flp-1* (333 bp, for AVK expression), *gcy-7* (1.2 kb, for ASEL expression), *tph-1* short (*tph-1s*, 158 bp, for NSM expression), *glr-3* (2.8 kb, for RIA expression), *tbh-1* (4.5 kb, for RIC expression), *glr-1* (3.1 kb), *flp-11* (2.5 kb, for RIS expression), *dat-1* (0.4 kb), *unc-17* (3.2 kb) and *unc-25* (1.8 kb), which were cloned into pPD95.75 or pPD95.77 vectors. The following cDNAs are used in the rescue experiments: *ric-4a* (0.6 kb), *ric-4b* (0.7 kb), *gpa-17* (1.2 kb), *glr-6* (2.6 kb), *jnk-1* (1.4 kb), *casy-1a* (3.0 kb), *casy-1b* (0.5 kb) and *bam-2* (3.0 kb). Transgenic animals were generated by microinjection-based germline transformation as described ^36^.

Briefly, purified DNA of constructs of interest at defined concentrations was mixed immediately before loading into a glass micropipette. Young adult hermaphrodites that contain less than five embryos are used for injection.

### RNA Isolation and Sequencing

For each experiment, about 15 and 50 plates (5.5 cm) of worms were used for RNA extraction for L4440 vector control and *atp-2* RNAi, respectively. Worms were washed with nuclease-free water followed by 10 minutes of vortex-mixing in GENEzol/TRIzol reagent (Geneaid) and 5 minutes of standing for bulk RNA extraction. Next, the sample was centrifuged for 1 minute, with the supernatant transferred to RNase-free tubes containing 99% ethanol (1 volume of GENEzol) and vortex-mixing. Subsequently, the sample was moved to the column followed by buffer washing. The column was then centrifuged for 3 minutes to dry the RNA samples, which were dissolved and eluted with 25 μl nuclease free water. Library construction was done according to the manufacture’s protocol (Illumina, San Diego, USA) using Agilent’s SureSelect XT HS2 mRNA preparation Kit for 150 base paired-end sequencing on Solexa platform (San Diego, USA). AMPure XP beads (Beckman Coulter, USA) were used for size selection.

### Bioinformatics Analysis of Transcriptomes

Raw sequences were obtained from the Illumina’s pipeline program bcl2fastq v2.20 and were expected to generate 60M (million reads or 9 Gb) per sample. Filtering of low-quality data and qualified reads was performed using Trimmomatic v0.36 ^37^.

Gene expression level was calculated as TPM (Transcript per Million). The reference genome (WBcel235(GCA_000002985.3)) and gene annotations were retrieved from Ensembl database. The DESeq2 ^38^ was employed to perform statistical analyses of gene expression profiles. Differentially expressed genes were defined based on published standards ^39^ with some modifications. We defined upregulated genes as a fold change ≥ 2 and downregulated ones as a fold change ≤ 0.5. The TPM threshold for genes with lower expression level in both up- and downregulated groups is set as 1. The filter criteria of differentially expression p-value < 0.05 was spared in the time course experiment as there was only one replica for each time point. Three replicas were performed for the memory formation / retrieval RNA-seq experiments. Gene ontology (GO) and enrichment analysis was performed using DAVID ^40^, “DAVID: https://david.ncifcrf.gov”

### Avoidance Assay with RNAi

Feeding RNAi was performed as described ^41^. Overnight-grown liquid culture of HT115 *E. coli* strain expressing the *atp-2* dsRNA or control (L4440) plasmid was diluted at 1:100 into 3 ml LB medium and incubated at 37°C for 3 h until OD600 reached 0.4. RNAi induction was achieved by adding 1 mM isopropyl β-D-1-thiogalactopyranoside (IPTG) and incubation at room temperature (25°C) for 1 h. Bacterial culture (100 μl) was seeded onto the center of the NGM plate containing 1 mM IPTG and ampicillin, and was allowed to grow overnight. Bleach-synchronized L1 larvae were carefully placed onto the center of the seeded RNAi plates. Bacterial avoidance was scored at 24, 36 and 48 h (L4440) or 54 h (*atp-2* RNAi) before worms were collected for RNA extraction with the avoidance index:

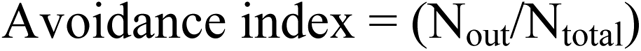

where N_out_ is the number of worms off-lawn and N_total_ is the number of total worms on the plate. As *atp-2* RNAi delays development, adult worms subjected to *atp-2* RNAi were collected at a later time point such that the developmental stages of the control and *atp-2* RNAi groups were comparable. In avoidance assays with transfer to RNAi-free plates, RNAi-treated animals were washed with M9, transferred to RNAi-free NGM plates seeded with HT115, and the avoidance index was calculated 20 h after transfer.

### Avoidance Assay with Antimycin A

OP50 *E. coli* in LB (100 μl, O.D. 0.4-0.6) was seeded at the center of the NGM plate to make a lawn of defined area the day before the avoidance assay. Antimycin A (Sigma-Aldrich) from the stock (2.5 mg/ml, in 99% EtOH) was diluted at 1:10 ratio with M9 buffer (0.25 mg/ml), and 100 µl of this antimycin A dilution was gently and evenly applied onto the bacteria lawn to reach the final concentration of 4.5 µM (2.5 μg/ml) in plate. Plates with lower concentrations of antimycin A were prepared similarly with adjusted amount of antimycin A. In the control group, 99 % ethanol (EtOH) was used in the place of the antimycin A stock. After the bacterial plate was completely dried in air, young adult animals were collected by washing with M9.

Worms were then gently transferred to the center of the assay plates. Bacterial avoidance was scored 6 h later or as indicated in individual experiments by the avoidance index described above.

### Bacterial Chemotaxis Assay

Bacterial chemotaxis assays were performed as described with some modifications ^13^. The 5.5-cm chemotaxis (CTX) plate (2 % agar, 5 mM potassium phosphate (pH 6.0), 1 mM CaCl_2_, 1 mM MgSO_4_) was divided into six equal longitudinal zones similar to that used for repulsive chemotaxis assays ^42^. One microliter of liquid OP50 (O.D. 0.4 - 0.6) and 1 µl of LB were spotted at the opposite sides of the plate, respectively. When these were completely dried, one microliter of 0.5 M sodium azide was applied to both spots to immobilize the worms that were attracted to reach OP50 or LB. D1 animals were treated with either ethanol or 1.125 µM antimycin (0.625 µg/ml) on small or full bacterial lawns as indicated in individual experiments for 3 h. This concentration of antimycin A was used because the standard 4.5 µM of antimycin A seemed to interfere with worm movements in the bacterial chemotaxis assay. Worms were then collected by washing with M9 for three times and transferred to NGM plates seeded with 200 μl HT115 *E. coli* for 15 h or as specified in individual experiments. The animals were then collected and washed, twice with the CTX wash buffer (5 mM potassium phosphate (pH 6.0), 1 mM CaCl_2_, 1 mM MgSO_4_) and once by water. Animals were transferred to the center of the chemotaxis plate and excessive liquid carefully removed by Kimwipes. The chemotaxis index (CI) was quantified 2 h later as

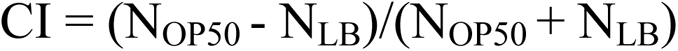

where N_OP50_ is the number of animals in the two zones close to OP50, and N_LB_ is the number of animals in the two zones close to LB.

### Heat Shock Rescue Experiments

Heat shock rescue experiments were performed using the bacterial chemotaxis assay as described above. To induce gene expression before memory formation, transgenic animals cultivated on regular OP50 plates were subjected to heat shock at 34°C for 1 h, followed by 1 h of recovery at 20°C. Animals were then washed three times with M9 and transferred to NGM plates seeded with OP50 *E. coli* containing 1.125 μM of antimycin for 3 h. To induce gene expression during memory retrieval, transgenic worms first underwent antimycin training (1.125 μM, 3 h). Animals were then transferred to antimycin-free HT115 *E. coli* plates for 15 h rest as described above. At the beginning of the 14^th^ hour of rest (i.e., 2 h before starting chemotaxis assay), worms were subjected to heat shock at 34°C for 1 h and allowed to recover for 1 h on the HT115 plates. Chemotaxis assays were then conducted as described above.

### Locomotion Assessment

For locomotion assessment, three to five well-fed young adult worms were picked to an unseeded plate to crawl for 3 to 5 minutes to shed the bacteria, and they were transferred again to an unseeded plate. Locomotion was videotaped for three minutes, and movement speed and reversal frequency were analyzed using WormLab (MBF Bioscience, Vermont, USA).

### Fluorescent Microscopy and the Coelomocyte Assays

D1 animals were immobilized in 1 % sodium azide on 5 % agar pad, and epifluorescent images were acquired by the AxioImager M2 system (Carl Zeiss).

Confocal projection images were acquired using the LSM700 Confocal Imaging System (Carl Zeiss). To quantify mRFP::CASY-1A signals in the coelomocytes of animals with the *Pcasy-1a::mRFP::CASY-1A::Venus* transgene, we identified the coelomocytes under DIC optics and confirmed with the BFP marker. Three endosomes with the highest mRFP signal intensity were quantified to obtain a mean mRFP value for each coelomocyte, followed by subtraction of background signals. To quantify mRFP::CASY-1A in animals exposed to bacterial odor, one agar plugs spotted with OP50 *E. coli* conditioned medium (grown to O.D. at 0.6) were placed on the lid of an unseeded plate, allowing the animals to be exposed to only volatile bacterial cues.

### Data Availability

The RNA sequencing data had been deposited in the NCBI Gene Expression Omnibus (GEO) under the accession number 243111.

## QUANTIFICATION AND STATISTICAL ANALYSIS

Statistical analyses, including two-way ANOVA followed by Bonferroni’s correction for multiple comparisons, multiple t-test, Mann-Whitney U test and Chi-Square test, were performed by Prism and described in the Figure Legends that apply, with sample sizes indicated.

## REFERENCES

1. Asok, A., Leroy, F., Rayman, J.B., and Kandel, E.R. (2019). Molecular Mechanisms of the Memory Trace. Trends Neurosci 42, 14–22.10.1016/j.tins.2018.10.005.

2. Modi, M.N., Shuai, Y., and Turner, G.C. (2020). The Drosophila Mushroom Body: From Architecture to Algorithm in a Learning Circuit. Annu Rev Neurosci 43, 465–484.10.1146/annurev-neuro-080317-0621333.

3. Lisman, J., Buzsáki, G., Eichenbaum, H., Nadel, L., Ranganath, C., and Redish, A.D. (2017). Viewpoints: how the hippocampus contributes to memory, navigation and cognition. Nat Neurosci 20, 1434–1447.10.1038/nn.4661.

4. Tonegawa, S., Morrissey, M.D., and Kitamura, T. (2018). The role of engram cells in the systems consolidation of memory. Nat Rev Neurosci 19, 485–498.10.1038/s41583-018-0031-2.

5. Davis, R.L. (2023). Learning and memory using Drosophila melanogaster: a focus on advances made in the fifth decade of research. Genetics 224.10.1093/genetics/iyad085.

6. Liu, H., and Zhang, Y. (2020). What can a worm learn in a bacteria-rich habitat? J Neurogenet 34, 369–377.10.1080/01677063.2020.1829614.

7. Roselli, C., Ramaswami, M., Boto, T., and Cervantes-Sandoval, I. (2021). The Making of Long-Lasting Memories: A Fruit Fly Perspective. Front Behav Neurosci 15, 662129.10.3389/fnbeh.2021.662129.

8. Jin, X., Pokala, N., and Bargmann, C.I. (2016). Distinct Circuits for the Formation and Retrieval of an Imprinted Olfactory Memory. Cell 164, 632–643.10.1016/j.cell.2016.01.007.

9. Kandel, E.R., Dudai, Y., and Mayford, M.R. (2014). The molecular and systems biology of memory. Cell 157, 163–186.10.1016/j.cell.2014.03.001.

10. Altarejos, J.Y., and Montminy, M. (2011). CREB and the CRTC co-activators: sensors for hormonal and metabolic signals. Nat Rev Mol Cell Biol 12, 141–151.10.1038/nrm3072.

11. Lakhina, V., Arey, R.N., Kaletsky, R., Kauffman, A., Stein, G., Keyes, W., Xu, D., and Murphy, C.T. (2015). Genome-wide functional analysis of CREB/long-term memory-dependent transcription reveals distinct basal and memory gene expression programs. Neuron 85, 330–345.10.1016/j.neuron.2014.12.029.

12. Melo, J.A., and Ruvkun, G. (2012). Inactivation of conserved C. elegans genes engages pathogen- and xenobiotic-associated defenses. Cell 149, 452–466.10.1016/j.cell.2012.02.050.

13. Chiang, Y.C., Liao, C.P., and Pan, C.L. (2022). A serotonergic circuit regulates aversive associative learning under mitochondrial stress in C. elegans. Proc Natl Acad Sci U S A 119, e2115533119.10.1073/pnas.2115533119.

14. Liao, C.P., Chiang, Y.C., Tam, W.H., Chen, Y.J., Chou, S.H., and Pan, C.L. (2022). Neurophysiological basis of stress-induced aversive memory in the nematode Caenorhabditis elegans. Curr Biol 32, 5309–5322.e5306.10.1016/j.cub.2022.11.012.

15. Kauffman, A.L., Ashraf, J.M., Corces-Zimmerman, M.R., Landis, J.N., and Murphy, C.T. (2010). Insulin signaling and dietary restriction differentially influence the decline of learning and memory with age. PLoS Biol 8, e1000372.10.1371/journal.pbio.1000372.

16. Jagannathan, S., Laughton, D.L., Critten, C.L., Skinner, T.M., Horoszok, L., and Wolstenholme, A.J. (1999). Ligand-gated chloride channel subunits encoded by the Haemonchus contortus and Ascaris suum orthologues of the Caenorhabditis elegans gbr-2 (avr-14) gene. Mol Biochem Parasitol 103, 129–140.10.1016/s0166-6851(99)00120-6.

17. Hou, Q., Gao, X., Zhang, X., Kong, L., Wang, X., Bian, W., Tu, Y., Jin, M., Zhao, G., Li, B., et al. (2004). SNAP-25 in hippocampal CA1 region is involved in memory consolidation. Eur J Neurosci 20, 1593–1603.10.1111/j.1460-9568.2004.03600.x.

18. Wang, C., Yang, B., Fang, D., Zeng, H., Chen, X., Peng, G., Cheng, Q., and Liang, G. (2018). The impact of SNAP25 on brain functional connectivity density and working memory in ADHD. Biol Psychol 138, 35–40.10.1016/j.biopsycho.2018.08.005.

19. Hill, E., Broadbent, I.D., Chothia, C., and Pettitt, J. (2001). Cadherin superfamily proteins in Caenorhabditis elegans and Drosophila melanogaster. J Mol Biol 305, 1011–1024.10.1006/jmbi.2000.4361.

20. Vogt, L., Schrimpf, S.P., Meskenaite, V., Frischknecht, R., Kinter, J., Leone, D.P., Ziegler, U., and Sonderegger, P. (2001). Calsyntenin-1, a proteolytically processed postsynaptic membrane protein with a cytoplasmic calcium-binding domain. Mol Cell Neurosci 17, 151–166.10.1006/mcne.2000.0937.

21. Hintsch, G., Zurlinden, A., Meskenaite, V., Steuble, M., Fink-Widmer, K., Kinter, J., and Sonderegger, P. (2002). The calsyntenins--a family of postsynaptic membrane proteins with distinct neuronal expression patterns. Mol Cell Neurosci 21, 393–409.10.1006/mcne.2002.1181.

22. Ikeda, D.D., Duan, Y., Matsuki, M., Kunitomo, H., Hutter, H., Hedgecock, E.M., and Iino, Y. (2008). CASY-1, an ortholog of calsyntenins/alcadeins, is essential for learning in Caenorhabditis elegans. Proc Natl Acad Sci U S A 105, 5260–5265.10.1073/pnas.0711894105.

23. 23. Hoerndli, F.J., Walser, M., Fröhli Hoier, E., de Quervain, D., Papassotiropoulos, A., and Hajnal, A. (2009). A conserved function of C. elegans CASY-1 calsyntenin in associative learning. PLoS One 4, e4880.10.1371/journal.pone.0004880.

24. Ding, C., Wu, Y., Dabas, H., and Hammarlund, M. (2022). Activation of the CaMKII-Sarm1-ASK1-p38 MAP kinase pathway protects against axon degeneration caused by loss of mitochondria. Elife 11.10.7554/eLife.73557.

25. Thapliyal, S., Vasudevan, A., Dong, Y., Bai, J., Koushika, S.P., and Babu, K. (2018). The C-terminal of CASY-1/Calsyntenin regulates GABAergic synaptic transmission at the Caenorhabditis elegans neuromuscular junction. PLoS Genet 14, e1007263.10.1371/journal.pgen.1007263.

26. Kim, B., and Emmons, S.W. (2017). Multiple conserved cell adhesion protein interactions mediate neural wiring of a sensory circuit in C. elegans. Elife 6.10.7554/eLife.29257.

27. Ohno, H., Kato, S., Naito, Y., Kunitomo, H., Tomioka, M., and Iino, Y. (2014). Role of synaptic phosphatidylinositol 3-kinase in a behavioral learning response in C. elegans. Science 345, 313–317.10.1126/science.1250709.

28. Katidou, M., Tavernarakis, N., and Karagogeos, D. (2013). The contactin RIG-6 mediates neuronal and non-neuronal cell migration in Caenorhabditis elegans. Dev Biol 373, 184–195.10.1016/j.ydbio.2012.10.027.

29. Lu, Z., Wang, Y., Chen, F., Tong, H., Reddy, M.V., Luo, L., Seshadrinathan, S., Zhang, L., Holthauzen, L.M., Craig, A.M., et al. (2014). Calsyntenin-3 molecular architecture and interaction with neurexin 1α. J Biol Chem 289, 34530–34542.10.1074/jbc.M114.606806.

30. Um, J.W., Pramanik, G., Ko, J.S., Song, M.Y., Lee, D., Kim, H., Park, K.S., Südhof, T.C., Tabuchi, K., and Ko, J. (2014). Calsyntenins function as synaptogenic adhesion molecules in concert with neurexins. Cell Rep 6, 1096–1109.10.1016/j.celrep.2014.02.010.

31. Colavita, A., and Tessier-Lavigne, M. (2003). A Neurexin-related protein, BAM-2, terminates axonal branches in C. elegans. Science 302, 293–296.10.1126/science.1089163.

32. Freytag, V., Probst, S., Hadziselimovic, N., Boglari, C., Hauser, Y., Peter, F., Gabor Fenyves, B., Milnik, A., Demougin, P., Vukojevic, V., et al. (2017). Genome-Wide Temporal Expression Profiling in Caenorhabditis elegans Identifies a Core Gene Set Related to Long-Term Memory. J Neurosci 37, 6661–6672.10.1523/jneurosci.3298-16.2017.

33. Hata, S., Fujishige, S., Araki, Y., Kato, N., Araseki, M., Nishimura, M., Hartmann, D., Saftig, P., Fahrenholz, F., Taniguchi, M., et al. (2009). Alcadein cleavages by amyloid beta-precursor protein (APP) alpha- and gamma-secretases generate small peptides, p3-Alcs, indicating Alzheimer disease-related gamma-secretase dysfunction. J Biol Chem 284, 36024–36033.10.1074/jbc.M109.057497.

34. 34. Araki, Y., Miyagi, N., Kato, N., Yoshida, T., Wada, S., Nishimura, M., Komano, H., Yamamoto, T., De Strooper, B., Yamamoto, K., et al. (2004). Coordinated metabolism of Alcadein and amyloid beta-protein precursor regulates FE65-dependent gene transactivation. J Biol Chem 279, 24343–24354.10.1074/jbc.M401925200.

35. Brenner, S. (1974). The genetics of Caenorhabditis elegans. Genetics 77, 71–94.10.1093/genetics/77.1.71.

36. Mello, C.C., Kramer, J.M., Stinchcomb, D., and Ambros, V. (1991). Efficient gene transfer in C.elegans: extrachromosomal maintenance and integration of transforming sequences. Embo j 10, 3959–3970.10.1002/j.1460-2075.1991.tb04966.x.

37. Bolger, A.M., Lohse, M., and Usadel, B. (2014). Trimmomatic: a flexible trimmer for Illumina sequence data. Bioinformatics 30, 2114–2120.10.1093/bioinformatics/btu170.

38. Love, M.I., Huber, W., and Anders, S. (2014). Moderated estimation of fold change and dispersion for RNA-seq data with DESeq2. Genome Biol 15, 550.10.1186/s13059-014-0550-8.

39. Ramsköld, D., Wang, E.T., Burge, C.B., and Sandberg, R. (2009). An abundance of ubiquitously expressed genes revealed by tissue transcriptome sequence data. PLoS Comput Biol 5, e1000598.10.1371/journal.pcbi.1000598.

40. Huang da, W., Sherman, B.T., and Lempicki, R.A. (2009). Systematic and integrative analysis of large gene lists using DAVID bioinformatics resources. Nat Protoc 4, 44–57.10.1038/nprot.2008.211.

41. Kamath, K., Oroudjev, E., and Jordan, M.A. (2010). Determination of microtubule dynamic instability in living cells. Methods Cell Biol 97, 1–14.10.1016/s0091-679x(10)97001-5.

42. Colbert, H.A., and Bargmann, C.I. (1995). Odorant-specific adaptation pathways generate olfactory plasticity in C. elegans. Neuron 14, 803–812.10.1016/0896-6273(95)90224-4.

